# Energy expenditure deficits drive obesity in a mouse model of Alström syndrome

**DOI:** 10.1101/2021.12.03.471145

**Authors:** Erin J Stephenson, Clint E Kinney, Amanda S Statyton, Joan C Han

**Author notes:** **Contact information** Corresponding author: Dr. Erin Stephenson, Mailing address: 555 31^st^ Street, SH-542L, Downers Grove IL, 60515, U.S.A. **Author contributions** E.J.S. and J.C.H. designed the studies. E.J.S., A.S.S. and C.E.K. carried out the experiments. E.J.S. analyzed and interpreted the data and prepared and revised the manuscript. J.C.H. provided feedback on the initial draft of the manuscript and its revision and all authors approved the submitted version of the manuscript.

## Abstract

Alström syndrome (AS) is a rare multi-system disorder for which early-onset childhood obesity is a cardinal feature. Like humans with AS, animal models with *Alms1* loss-of-function mutations develop obesity, supporting the notion that ALMS1/Alms1 is required for the regulatory control of energy balance across species. This study aimed to determine which component(s) of energy balance are reliant on Alms1. Here, we performed comprehensive energy balance phenotyping *Alms1*^*tvrm102*^ mice at both eight- and eighteen-weeks-of-age. We found that adiposity gains occurred early and rapidly in *Alms1*^*tvrm102*^ male mice but much later in females. Rapid increases in body fat in males was due to a marked reduction in energy expenditure (EE) during early life and not due to any genotype-specific increases in energy intake under chow conditions. Energy intake did increase in a genotype-specific manner when mice were provided a high-fat-diet, exacerbating the effects of reduced EE on obesity progression. The EE deficit observed in male *Alms1*^*tvrm102*^ mice did not persist as mice aged, suggesting loss of Alms1 either causes a developmental delay in the mechanisms controlling early life EE, or that activation of compensatory mechanisms occurs after obesity is established. Future studies will determine how ALMS1/Alms1 modulates EE and how sex moderates this process.

## Introduction

Primary cilia are membranous structures that protrude from the surface of all quiescent cells and many terminally differentiated cells [1]. They are dynamic organelles that sense changes in the extracellular environment and communicate these changes to the cell [1, 2]. The localization and trafficking of receptors, ion channels and other proteins along the length of the cilium is essential for maintaining coordination of intracellular functions [1-4], and syndromes associated with defective cilia frequently manifest as a collection of metabolic defects and multi-organ dysfunction [5].

*ALMS1* (Chr-2q-3) encodes a 461-kDa protein located in the centrosomes and basal body of primary cilia [6, 7]. Although ALMS1 function is not completely understood [8, 9], humans with recessively inherited loss-of-function mutations manifest a ciliopathy known as Alström syndrome (AS) [8-10]. AS is an extremely rare multi-system disorder characterized by early-onset childhood obesity, cone-rod retinal dystrophy, and sensorineural hearing loss [8-11]. Dilated cardiomyopathy, type-2 diabetes, and progressive dysfunction of other organ systems (pulmonary, hepatic, and renal) are also frequently observed [8, 9, 11].

Mice with *Alms1* mutations have normal ciliary formation [12, 13], but cilia progressively deteriorate, leading to a reduction in the total number of functional primary cilia with advancing age [13-15]. This phenomenon appears to occur randomly [15] and across multiple tissues [13-15], with especially rapid deterioration reported in the hypothalamic region of the brain [13]. Like humans with AS, other animals with *Alms1* loss-of-function mutations develop obesity and hyperinsulinemia [12, 13, 15-18], suggesting ALMS1/Alms1 and long-term ciliary maintenance are required for effective regulatory control of energy balance.

Despite the observation that animals with *Alms1* loss-of-function mutations gain more bodyweight than their wild-type (WT) littermates [12, 15-17, 19-21], a comprehensive energy balance phenotype has not been established for any of the models reported to date (Table 1). Increased adiposity [12, 15, 16, 20, 21] and increased food intake [16, 21] have been observed, and progressive hyperglycemia [12, 16, 20, 21] and hyperinsulinemia [12, 20] have also been reported. Since pair-feeding only partially attenuates the obesity phenotype in *Alms1-*null mice [21], we hypothesized that obesity due to loss of Alms1 function may be driven via mechanisms of reduced energy expenditure (EE). Here, we have characterized the energy balance phenotype of the Alms1^*Tvrm102*^ mouse model of AS. We report sex-specific EE deficits in the absence of hyperphagia during the early stages of obesity development.

**Table 1:** Overview of mouse models with systemic *Alms1* mutations and their metabolic abnormalities

| Model | Mutation | Background | Metabolic observations compared to WT littermates | Citation(s) |
| --- | --- | --- | --- | --- |
| *foz/foz* fed a chow diet | Spontaneous 11 bp deletion in exon 8 causing a frameshift mutation that generates a premature stop | NOD (Non Obese Diabetic) | • ↑ Body mass index<br>• ↑ Adiposity<br>• ↑ Lean body mass<br>• ↑ Body length (females only)<br>• ↑ Daily food intake | [16, 21] |
|  | codon |  | • Hepatic steatosis<br>• Hyperinsulinemia<br>• Hyperglycemia<br>• ↓ Glucose tolerance<br>• Enlarged pancreatic islets<br>• ↑ Serum cholesterol (total)<br>• ↔ $BW^{0.66}$ normalized $VO_2$ (males)<br>• ↔ Habitual physical activity (males) |  |
| *foz/foz* fed a HFD (males only) | Spontaneous 11 bp deletion in exon 8 causing a frameshift mutation that generates a premature stop codon | NOD (Non Obese Diabetic) | • > ↑ Bodyweight<br>• > ↑ Adiposity<br>• > ↑ Adipocyte hypertrophy<br>• > ↑ Daily food intake<br>• Hepatic steatosis<br>• Hyperglycemia<br>• ↓ Glucose tolerance<br>• ↓ Insulin responsiveness<br>• ↔ Body temperature<br>• ↓ $BW^{0.66}$ normalized $VO_2$<br>• ↔ Habitual physical activity | [21] |
| Gt(XH152) | Gene trap – intron 13 | C57BL6/J x 129P2Ola | • ↑ Bodyweight<br>• ↑ Adiposity<br>• Adipocyte hypertrophy<br>• Hepatic steatosis<br>• Hyperinsulinemia<br>• Hyperglycemia<br>• ↓ Glucose tolerance<br>• Enlarged pancreatic islets<br>• ↑ Serum leptin<br>• ↑ Serum cholesterol (total)<br>• ↑ Serum ALT (late)<br>• ↔ Serum triglycerides | [12, 20] |
| L2131X/L2131X | ENU-generated nonsense mutation in exon 10 TTA>TAA that generates a premature stop codon | C57BL6/J x NOD | • ↑ Bodyweight<br>• ↑ Adiposity<br>• Adipocyte hypertrophy<br>• Hepatic steatosis<br>• Hyperinsulinemia<br>• Euglycemia<br>• ↑ Serum leptin<br>• ↑ Serum cholesterol (total & HDL)<br>• ↑ Serum triglycerides | [15] |
| *tvrm102* | ENU-generated splice mutation in exon 6 T>C generating a premature stop codon | C57BL6/J | • ↑ Bodyweight | [19] |
| *flin/flin* (3-month-old males) | Frt-lacZ-LoxP-Neo-Frt-LoxP sequence inserted between exon 6 and exon 7 | C57BL6/N | • ↑ Bodyweight<br>• Adipocyte hypertrophy<br>• ↓ Glucose tolerance<br>• ↓ Insulin responsiveness<br>• Metabolic defects corrected with *Adipoq*-Cre driven *Alms1* reactivation | [17] |

## Methods

### Animals

Experimental animals were generated by heterozygous cross of *Alms1*^*tvrm102*^ mice (C57BL/6J-Alms1^*tvrm102/PjnMmjax*^, MMRRC Stock No: 43589-JAX, The Jackson Laboratory). This strain originated as an ENU-induced T>C point mutation in the splice donor site of exon 6, which unmasks a cryptic splice site 120 bases downstream [19]. Mice were genotyped using a custom Taqman SNP assay (Applied Biosystems #4332075, ANMFW9M). Homozygous mice (*Alms1*-/-, n=44) and WT litter mates (WT, n=52) were studied. Mice were housed at ∼22°C in ventilated cages with cellulose bedding, free access to chow (Envigo #7012), and an automated water supply. To determine whether food palatability influenced intake, one cohort was given free access to a high-fat, sucrose-containing diet for two-weeks (HFD; Research Diets #D12451), from 15-weeks-of-age. In this cohort, food intake was measured daily for four-weeks (one-week of regular chow from 14-weeks-of-age, two-weeks of HFD, and another week of chow; Figure 6A). The UTHSC IACUC approved all procedures.

### Metabolic phenotyping

From four-weeks-of-age, mixed genotype, group-housed mice were weighed, and body composition measured weekly by magnetic resonance (EchoMRI; Figure 1A). Body length was determined (distance from nose to base of tail) in isoflurane-anesthetized mice at nine- and 19-weeks-of-age. At eight-or 18-weeks-of-age, chow-fed mice were individually housed for two days before being moved to a high-resolution home cage-style Comprehensive Laboratory Animal Monitoring System (Columbus Instruments) which simultaneously records respiratory gases via open-circuit indirect calorimetry, physical activity via infrared beam breaks, and food intake via load cell-linked hanging feeder baskets. EE was calculated from VO_2_ and VCO_2_ using the Weir equation [22]. Resting EE was calculated from EE measured during ZT 0-3 when mice were not actively moving/eating. Non-resting EE was calculated as the difference between total EE and resting EE. The thermic effect of food (TEF) was determined from the energy content of the diet and amount of food consumed, with the difference between non-resting EE and TEF considered activity EE [23]. Energy balance was calculated as the difference between energy intake and EE, whereas metabolic efficiency was calculated as the percent of energy intake stored in fat and lean tissues [24]. Rates of fat and carbohydrate oxidation were calculated from VO_2_ and VCO_2_ using previously reported equations [25] with the assumption protein oxidation was negligible. Total activity was calculated as the combined number of beam breaks along the X- and Y- axes, whereas ambulatory activity was calculated as the combined number of consecutive X- and Y-axes beam breaks occurring in series. The difference in beam breaks between ambulatory and total activity was considered stereotypic activity (grooming, rearing and other non-ambulatory movement). Food intake was determined from individual feeding bouts from which cumulative intake was calculated. Data for each mouse was collected over 7-days, under ambient (22°C, 2-3-days) and thermoneutral conditions (28°C, 2-3-days; to mimic more closely the metabolic conditions of humans) [26]. After discarding initial light and dark photoperiods to account for acclimation, data were binned by hour and hourly mean values (indirect calorimetry data) or summed values (activity and food intake) over each 24-h measurement period were used to determine hourly values for each mouse. Data were analyzed using mixed linear models with bodyweight, body fat and fat-free mass (FFM) included as random effect variables. Total daily values were also calculated and analyzed by ANVOCA with fat and FFM as covariates. Figures show non-normalized group mean data ± standard error.

**Figure 1:**
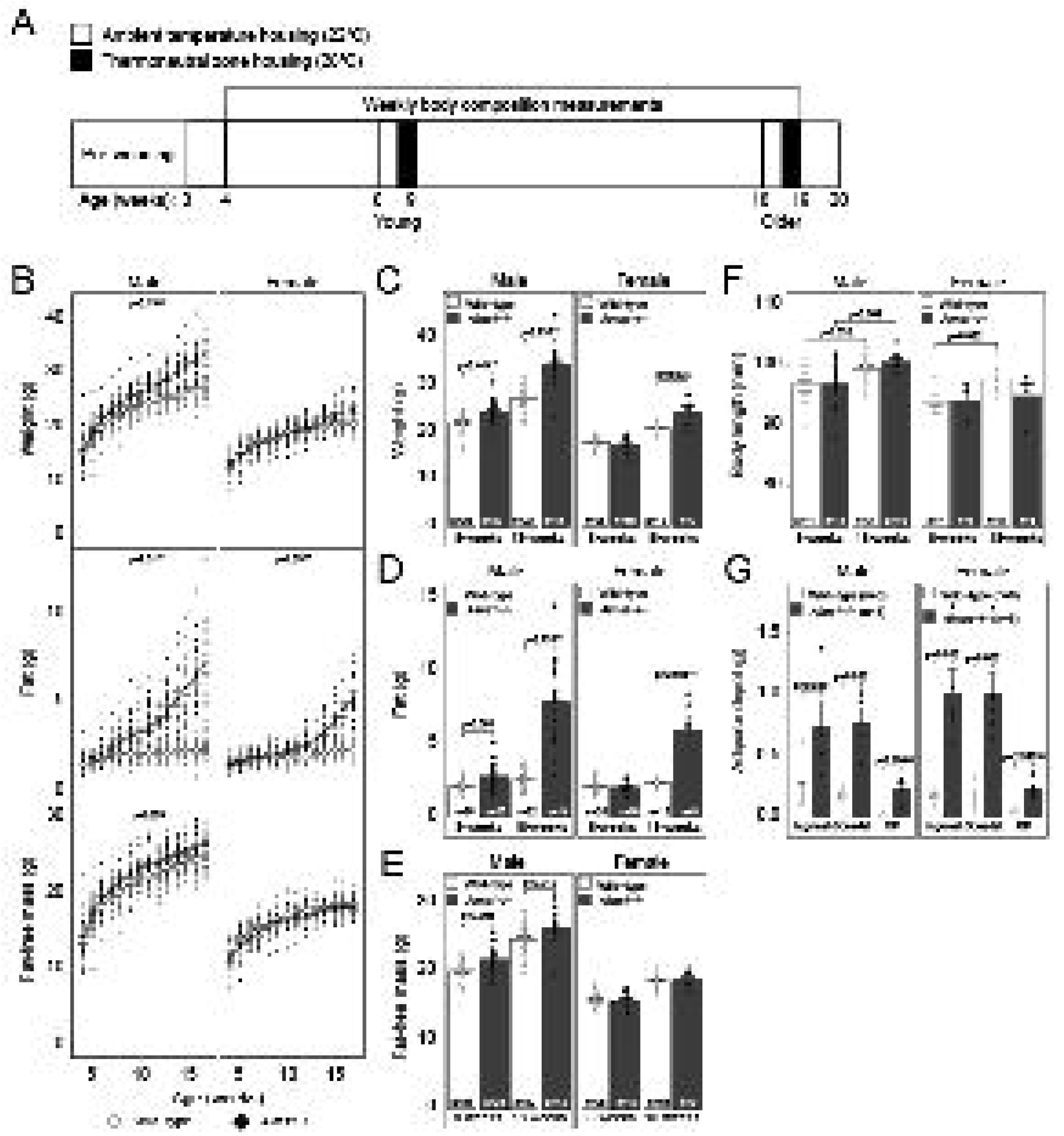
Body composition of *Alms1-*/- and WT mice. A. Overview of experimental design. B. Body composition of *Alms1-*/- and WT male (n=16-25 and n=21-24, respectively) and female mice (n=11-15 and n=19-22, respectively) from 4 weeks of age onward. C. Bodyweight at 8- and 18-weeks-of-age. D. Adiposity at 8- and 18-weeks-of-age. E. Fat-free-mass at 8- and 18-weeks-of-age. F. Body length at 9- and 19-weeks-of-age. G. Mass of adipose depots 20-weeks-of-age. Values displayed are mean ± se with individual data overlayed. Open circles/bars denote WT, closed circles/bars denote *Alms1*-/-.

### Statistics

Data are presented as group means ± standard error unless otherwise stated. Longitudinal data were analyzed by mixed linear models with likelihood-ratio tests. All other data were assessed for homoscedasticity (Levene’s test) and normality (Shapiro-Wilk test) before being compared as follows: normal data were analyzed using two-way ANOVA with Tukey post-hoc analysis; any multi-factorial non-normal data were analyzed using Scheirer-Ray-Hare tests with post hoc Wilcoxon Rank Sum tests adjusted for multiple comparisons using the Benjamini and Hochburg correction [27]. Pairwise comparisons were made using Student’s t-tests, Welch’s t-tests or Wilcoxon Rank Sum tests. Data wrangling and analyses were performed in RStudio using R v4.1.1 and statistical packages lme4 [28] (linear models), car [29] (Levene’s tests), rcompanion [30] (Scheirer-Ray-Hare tests), rstatix [31] and emmeans [32] (ANCOVA/estimated marginal means). Differences between groups are discussed as significant where p-values are <0.05.

## Results

### Adiposity gains occur early in Alms1-/-males but later in females

To characterize the development and progression of obesity resulting from loss of Alms1, we established body composition trajectories of male and female *Alms1*-/-mice starting from four-weeks-of-age (Figure 1B). Compared to WT, *Alms1*-/-mice progressively gained more adiposity, with obesity occurring earlier in males and to a much greater magnitude than in females. Compared to WT, *Alms1*-/-males had 20.4% more adiposity at five-weeks-of-age (p=0.004), 6.9% greater bodyweight (p=0.012), and by six-weeks-of-age, 5.1% more FFM (p=0.031). By 18-weeks-of-age (Figures 1C-E), *Alms1*-/-males were 24.0% heavier (p<0.001) with 195.3% more adiposity (p<0.001) and 6.7% more FFM than WT (p=0.004). In contrast, *Alms1*-/-females weighed 7.5% less (p=0.047) than WT at five-weeks-of-age. Increased adiposity was not observed in *Alms1*-/-females until 10-weeks-of-age (17.5%, p=0.009), whereas total body weight did not diverge from WT until 16-weeks-of-age (10.0%, p=0.021), at which point adiposity was 96.3% greater than WT (p<0.001). By 18-weeks-of-age, *Alms1*-/-females were 11.9% heavier than WT, with 119.7% more adiposity (p<0.001; Figures 1C-D). No differences in FFM were observed between genotypes in female mice (Figures 1B and 1E). Although an effect of age was observed for body length in both sexes (p<0.001 and p=0.01), differences in body composition were not driven by genotype-dependent differences in body length in male mice. In contrast, genotype influenced female body length (p=0.052; Figure 1F), with WT growing 4% longer between nine and nineteen-weeks-of-age (p=0.022) during which time the length of *Alms1*-/- females remained reasonably stable (p=0.978). Increases in adiposity in both sexes occurred similarly across inguinal, gonadal, and retroperitoneal adipose depots (Figure 1G).

### Young Alms1-/- mice are less active than WT and males have lower EE

To identify which determinants of energy balance contribute to the increased adiposity we observed in *Alms1*-/- mice, we measured EE while simultaneously measuring physical activity and energy intake (Figures 1A, 2-5). We initially performed these experiments in eight-week-old mice to limit the confounding influence of genotype-specific differences in bodyweight in older mice. Under ambient conditions (22°C, Figure 2A-C), compared to WT, 8-week-old *Alms1-/-* males had reduced EE (p=0.032; Figure 2A), a finding consistent across both light (p=0.052) and dark (p=0.024) photoperiods. When body composition was taken into account, the genotype effect on EE persisted (p=3.0^-04^), with *Alms1*-/- having reduced EE in both the light (p=4.6^-04^) and dark photoperiods (p=3.0^-04^). Reduced EE was associated with 58.4% lower activity EE (p=0.026; Table 2). Resting EE was not different (Figure 2B, shaded bars), neither was the thermic effect of food (Table 2). Genotype explained a 2.09 kCal/d difference in daily EE in male *Alms1*-/- mice compared to WT when body fat and FFM were matched (2.13 and 20.06 g, respectively, p=0.001; Figure 2C). Lower EE was due, in part, to *Alms1*-/- mice being 22.0% less ambulatory (Figures 2D-E, p=0.043 and p=0.030), especially during the dark photoperiod (Figure 2D, p=0.028). Lower total-EE (16.3%, p=0.028) and reduced ambulation (25.1%, p=0.009) persisted in male *Alms1*-/- under thermoneutral conditions (28°C; Supplementary Figure 1A) indicating EE deficits in male *Alms1*-/- were not a consequence of genotype-specific responses to thermal stress caused by housing mice at temperatures considered ambient for humans but cold for mice [26]. Given EE was lower in male *Alms1-/-*, energy balance tended to be higher (p=0.084; Figure 2F), despite food intake being similar to WT (Figures 2G-H). In contrast, none of the EE parameters we measured differed between genotypes in female mice under either housing temperature. Due to light photoperiod ambulation being lower in *Alms1*-/- females (p=0.031; Figure 2D), there was a tendency for total ambulation to be lower in *Alms1*-/- compared to WT at 22°C, although this was not significant (Figure 2D-E; p=0.073 and p=0.090) unless body composition was included in the model (p=0.045; Figure 2D). Given both EE and energy intake were similar between *Alms1*-/- and WT females, energy balance was also similar (p=0.183; Figure 2F). No genotype effect was observed for metabolic efficiency for either sex (Figure 2I).

**Table 2:** Average daily energy expenditure values for chow-fed mice

| Age | Sex | Genotype | Total EE |  | Resting EE |  | Food EE |  | Activity EE |  |
| --- | --- | --- | --- | --- | --- | --- | --- | --- | --- | --- |
|  |  |  | kCal/d | N | kCal/d | N | kCal/d | N | kCal/d | N |
| 8-9 weeks | Male | Wild-type | 10.1 ± 0.5 | 12 | 7.7 ± 0.5 | 8 | 1.2 ± 0.1 | 12 | 0.8 ± 0.1 | 7 |
|  |  | *Alms1*<sup>-/-</sup> | 8.8 ± 0.4* | 9 | 7.0 ± 0.3 | 8 | 1.3 ± 0.1 | 9 | 0.4 ± 0.1* | 7 |
|  | Female | Wild-type | 9.0 ± 0.5 | 12 | 6.8 ± 0.4 | 6 | 1.2 ± 0.05 | 12 | 0.7 ± 0.2 | 6 |
|  |  | *Alms1*<sup>-/-</sup> | 9.7 ± 0.5 | 10 | 7.6 ± 0.3 | 9 | 1.2 ± 0.04 | 10 | 0.9 ± 0.3 | 8 |
| **18-19 weeks** | Male | Wild-type | 11.0 ± 0.3 | 22 | 8.7 ± 0.2 | 19 | 1.9 ± 0.1 | 10 | 0.8 ± 0.1 | 6 |
|  |  | *Alms1*<sup>-/-</sup> | 12.0 ± 0.3 | 18 | 10.2 ± 0.3* | 18 | 1.5 ± 0.1 | 8 | 0.6 ± 0.1 | 6 |
|  | Female | Wild-type | 10.5 ± 0.3 | 17 | 8.4 ± 0.2 | 16 | 1.6 ± 0.1 | 6 | 1.1 ± 0.2 | 5 |
|  |  | *Alms1*<sup>-/-</sup> | 10.6 ± 0.5 | 10 | 8.9 ± 0.5 | 10 | 1.5 ± 0.1 | 4 | 0.7 ± 0.1 | 3 |

Values are mean kCal/d ± se. \*p<0.05 different from same-sex, same age wild-type littermates by ANCOVA with body fat and fat-free mass as covariates.

**Figure 2:**
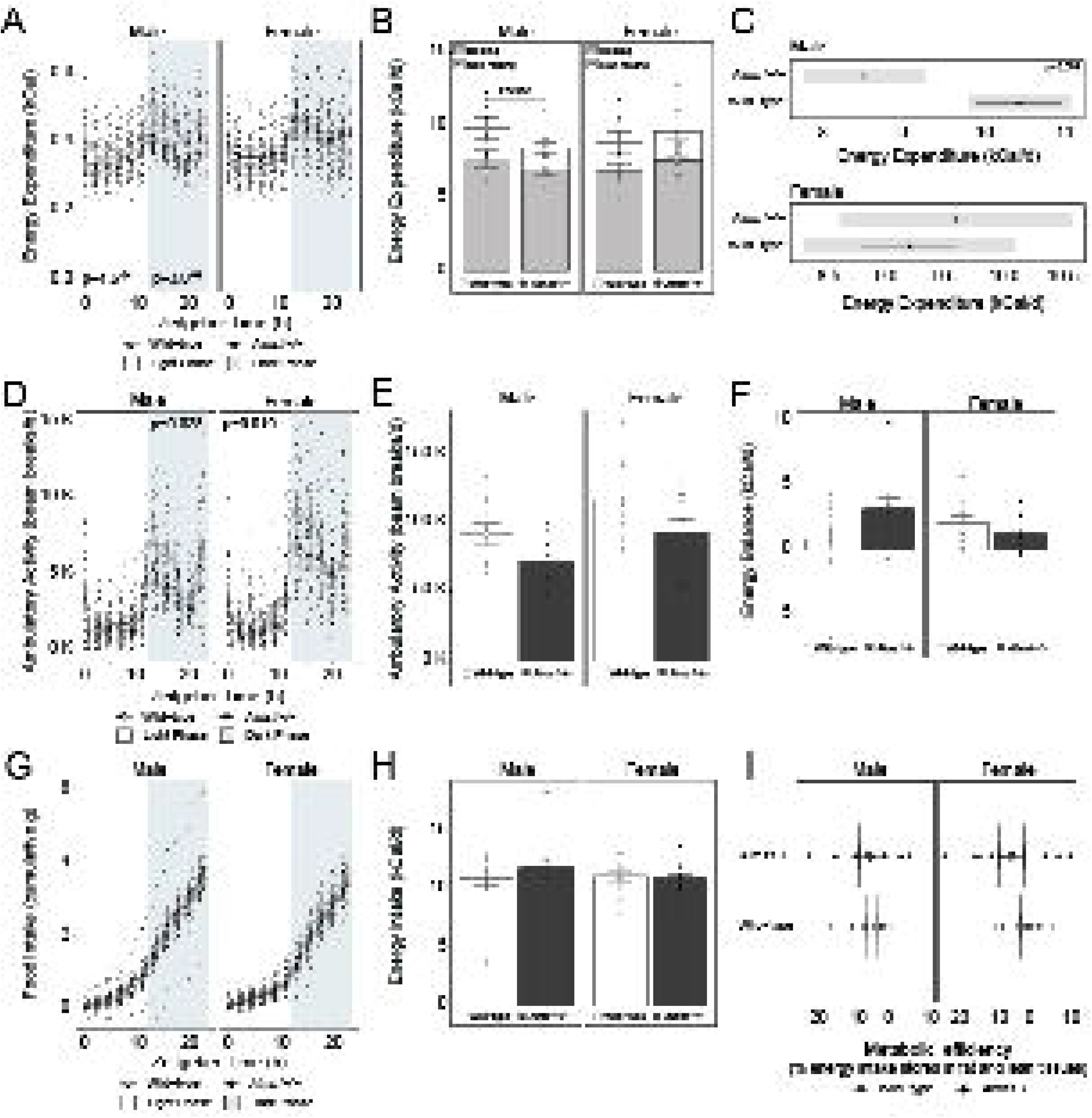
Energy balance phenotype of *Alms1-*/- and WT mice between 8-9-weeks-of-age. A. Hourly energy expenditure of young *Alms1*-/- and WT male (n=8 and n=11, respectively) and female mice (n=5 and n=9, respectively) measured at 22°C. B. Resting (open bars, closed circles) and nonresting (shaded bars, open circles) energy expenditure. C. Estimated marginal means for total daily energy expenditure for male and female mice assuming matched fat and fat-free mass. D. Ambulation across the day and E. total daily ambulatory activity. F. Energy balance. G. Cumulative and H. total daily energy intake. I. Metabolic efficiency. Data presented are group means ± se with individual data points overlayed. Open circles/bars denote WT, closed circles/bars denote *Alms1*-/-. Grey shading in A, D and G denotes the dark photoperiod. Grey shading in C denotes 95% confidence intervals.

### Young male Alms1-/- mice have altered substrate oxidation

To determine whether Alms1 loss affected the capacity to oxidize different energy substrates, we leveraged the indirect calorimetry data to calculate rates of lipid and carbohydrate oxidation (Figure 3). In male *Alms1*-/-, absolute lipid oxidation was not different from WT (p=0.189) but was lower during the light photoperiod (p=0.039). However, when body composition was considered, lipid oxidation was suppressed in *Alms1*-/- (p=0.9.4^-04^; Figures 3A-B), with genotype explaining an 84.4 mg/d difference in lipid oxidation (p=0.003; Figure 3C). Similarly, rates of carbohydrate oxidation were also lower in *Alms1*-/- males compared to WT (Figures 3D-F) both by absolute values (p=0.012) and when body composition was considered (Figure 3D-E; p=0.002 and p=0.005). These observations were driven by lower rates of carbohydrate oxidation during both light and dark photoperiods (Figure 3D; p=0.025 and p=0.003, respectively). Genotype explained a 348.8 mg/d l difference in carbohydrate oxidation (p=0.005; Figure 3F). Differences in lipid oxidation persisted under thermoneutral conditions (p=0.011; Supplementary Figure 1A). In contrast, differences in lipid oxidation were not significant in female mice (Figures 3A-B) despite *Alms1-/-* females having higher rates of lipid oxidation during the dark photoperiod compared to WT (p=0.039; Figure 3A). Inclusion of body composition in the model did not change these findings, whereas housing mice under thermoneutral conditions also made no difference. Rates of carbohydrate oxidation were similar between genotypes in female mice (Figures 3D-F), regardless of model design or housing temperature.

**Figure 3:**
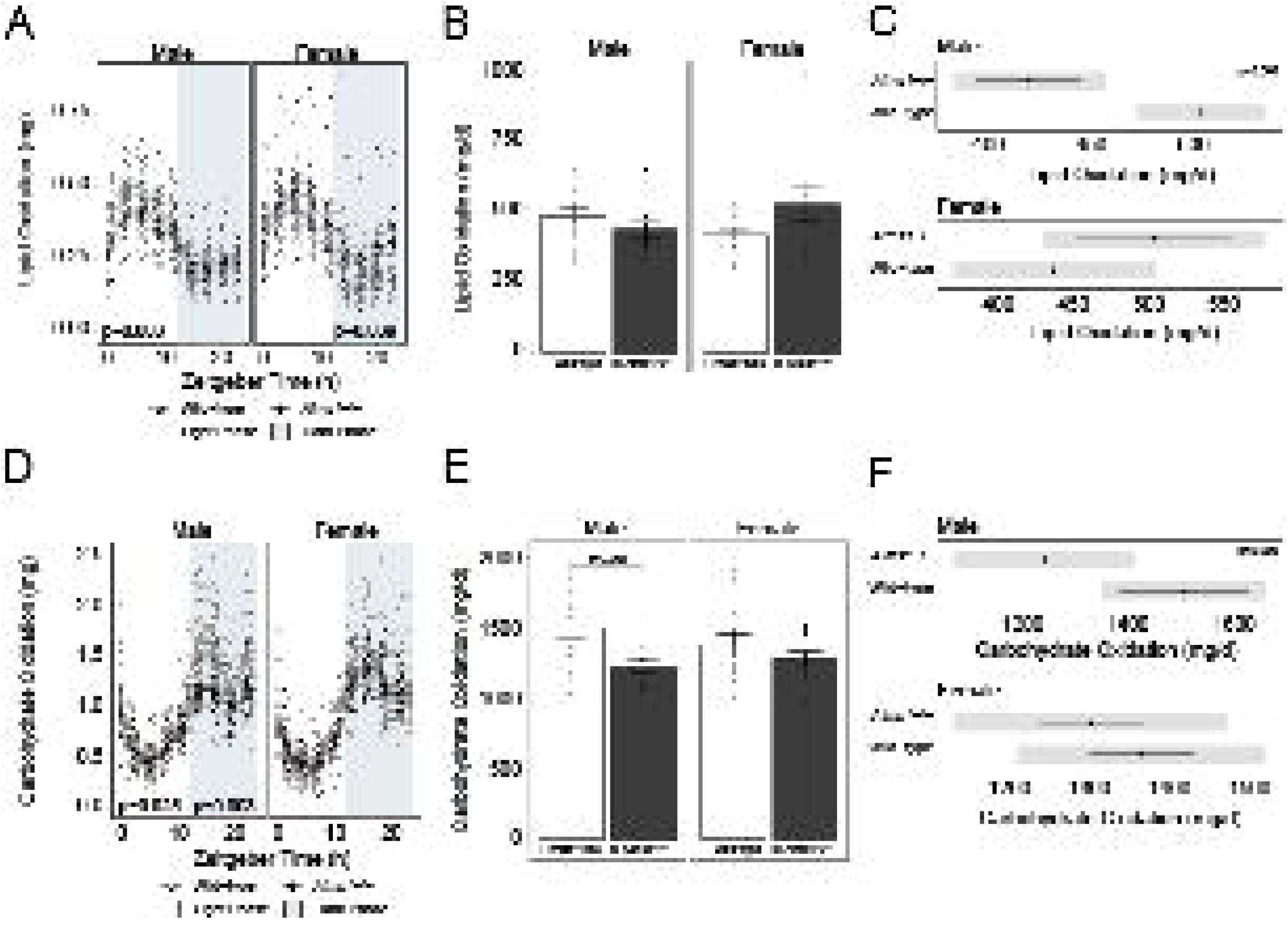
Rates of substrate oxidation in *Alms1-*/- and WT mice between 8-9-weeks-of-age.

A. Hourly lipid oxidation of young *Alms1*-/- and WT male (n=8 and n=11, respectively) and female mice (n=5 and n=9, respectively) measured at 22°C. B. Total daily lipid oxidation C. Estimated marginal means for daily lipid oxidation for male and female mice assuming matched fat and fat-free mass. D. Hourly carbohydrate oxidation of young *Alms1*-/- and WT male (n=8 and n=11, respectively) and female mice (n=5 and n=9, respectively) measured at 22°C. E. Total daily carbohydrate oxidation F. Estimated marginal means for daily carbohydrate oxidation for male and female mice assuming matched fat and fat-free mass. Data presented are group means ± se with individual data points overlayed. Open circles/bars denote WT, closed circles/bars denote *Alms1*-/-. Grey shading in A and D denotes the dark photoperiod. Grey shading in C and F denotes 95% confidence intervals.

### Aging attenuates EE deficits observed in younger Alms1-/- males

Since differences body composition did not manifest in *Alms1-/-* females until a later stage and no deficit in EE was observed in 8-week-old females, we repeated indirect calorimetry experiments in a cohort of older mice to determine if females might acquire a similar energy balance phenotype to male mice as they age (Figures 4A-I). Similar to our findings in younger mice, no differences in any of the EE parameters were observed in *Alms1-/-* females compared to WT (Figures 4A-C, Table 2) despite *Alms1-/-* females having increased adiposity by this age. Also like our observations in younger females, total ambulation tended to be reduced in *Alms1*-/- but this was not significant (Figure 4D), even when body composition was factored into the model (Figure 4D-E). Energy balance was not different between genotypes (Figure 4F) as neither EE or food intake differed (Figures 4G-H), and despite older female *Alms1*-/- mice having increased adiposity compared to WT, no difference in metabolic efficiency was observed (Figure 4I). In contrast to our observations in younger male mice, EE in older *Alms1-/-* males tended to be increased compared to WT (p=0.063; Figure 4A), although when body composition was considered, EE was not different between groups. When total EE was separated into resting and non-resting components (Figure 4B, Table 2), we observed 16.8% greater resting EE in older *Alms1*-/- males compared to WT (p=0.043) but no difference in non-resting EE. Activity differences present in younger mice were also no longer present in older males (Figures 4D-E). Older *Alms1*-/- males were also closer to neutral energy balance than WT (p=0.029; Figure 4F), a finding associated with a tendency for *Alms1*-/- mice to consume less food (Figures 4G-F). Unlike younger mice, metabolic efficiency was 1298.2% lower in older *Alms1*-/- males compared to WT (p=0.025; Figure 4I).

**Figure 4:**
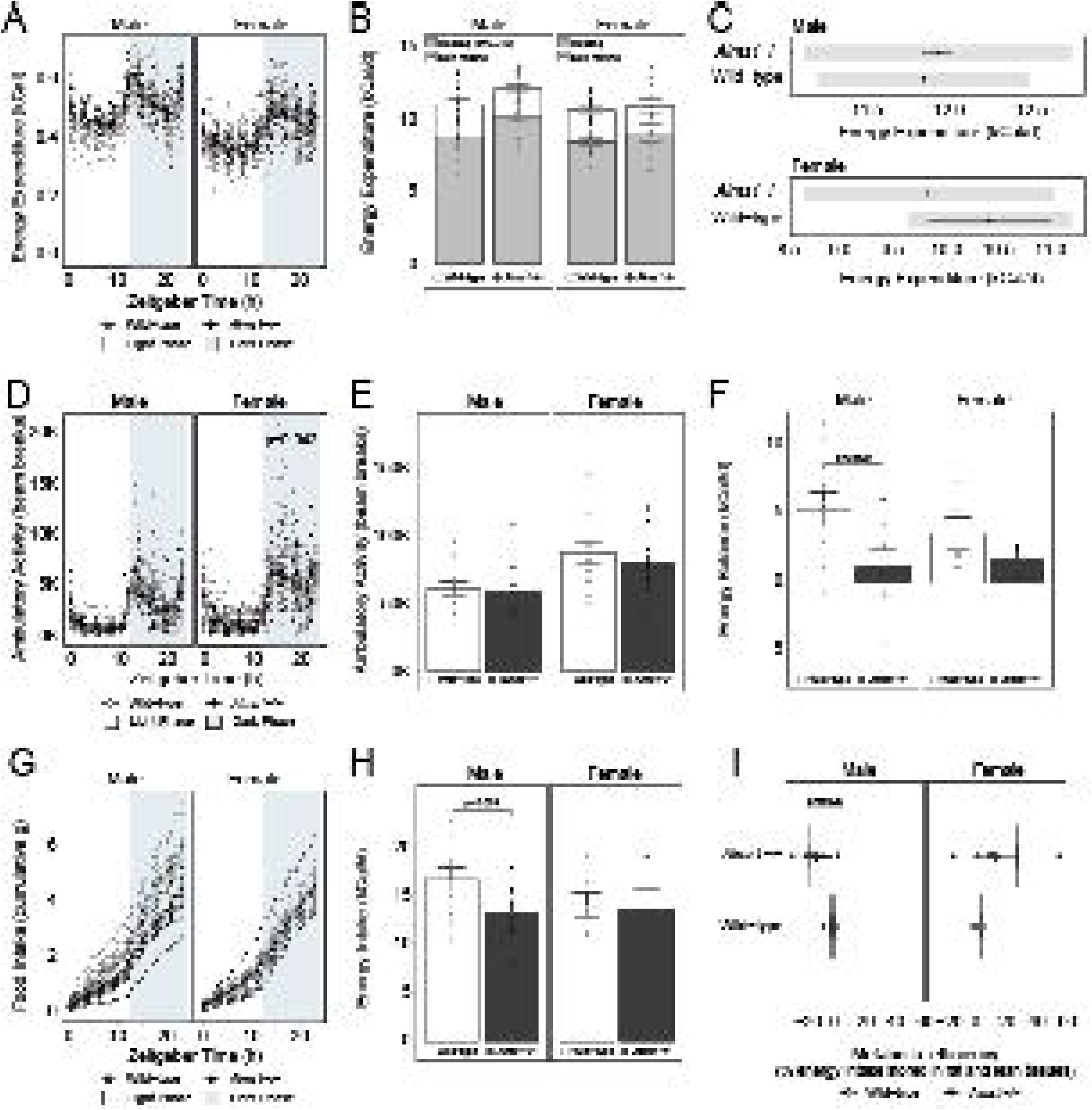
Energy balance phenotype of *Alms1-*/- and WT mice between 18-19-weeks-of-age. A. Daily energy expenditure of older *Alms1*-/- and WT male (n=8 and n=11, respectively) and female mice (n=10 and n=17, respectively) measured at 22°C. B. Resting (open bars; closed circles) and nonresting (shaded bars, open circles) energy expenditure. C. Estimated marginal means for total daily energy expenditure for male and female mice assuming matched fat and fat-free mass. D. Ambulation across the day and E. total daily ambulatory activity. F. Energy balance. G. Cumulative and H. total daily energy intake. I. Metabolic efficiency. Data presented are group means ± se with individual data points overlayed. Open circles/bars denote WT, closed circles/bars denote *Alms1*-/-. Grey shading in A, D and G denotes the dark photoperiod. Grey shading in C denotes 95% confidence intervals.

### Altered substrate oxidation can be explained by body composition in older Alms1-/- mice

Unlike our observations in younger mice, when comparing absolute lipid oxidation values, we observed increased lipid oxidation in both female and male *Alms1*-/- compared to their WT counterparts (Figure 5A-B; p=0.047 and p=0.004). However, no differences in lipid oxidation were detected after considering body composition in the comparisons (Figure 5C). Similarly, carbohydrate oxidation was decreased in female *Alms1*-/- compared to WT when absolute values were compared (p=0.042; Figure 5D-E), but when body composition was considered, rates of carbohydrate oxidation were similar to WT (Figure 5F). Carbohydrate oxidation did not differ in older male mice whether body composition was included in the model or not. Similar observations were made for all parameters in both sexes when older mice were housed under thermoneutral conditions (Supplementary Figure 1B); the only exception being that male *Alms1*-/- mice had lower fat oxidation during the dark photoperiod (p=0.042) and tended to have increased carbohydrate oxidation during the light photoperiod (p=0.076).

**Figure 5:**
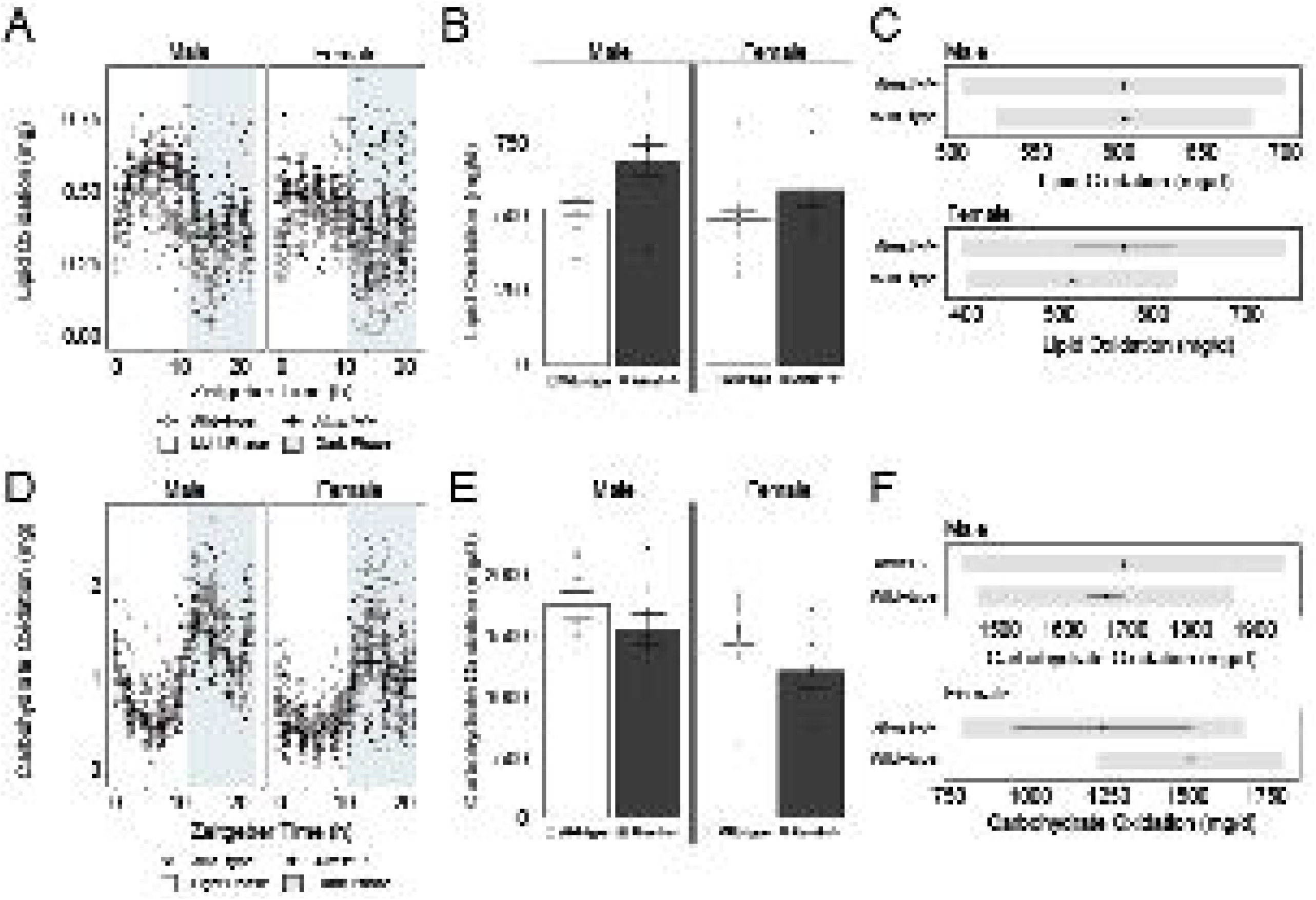
Rates of substrate oxidation in *Alms1-*/- and WT mice between 18-19-weeks-of-age A. Hourly lipid oxidation of young *Alms1*-/- and WT male (n=8 and n=11, respectively) and female mice (n=10 and n=17, respectively) measured at 22°C. B. Total daily lipid oxidation C. Estimated marginal means for daily lipid oxidation for male and female mice assuming matched fat and fat-free mass. D. Hourly carbohydrate oxidation of young *Alms1*-/- and WT male (n=8 and n=11, respectively) and female mice (n=5 and n=9, respectively) measured at 22°C. E. Total daily carbohydrate oxidation F. Estimated marginal means for daily carbohydrate oxidation for male and female mice assuming matched fat and fat-free mass. Data presented are group means ± se with individual data points overlayed. Open circles/bars denote WT, closed circles/bars denote *Alms1*-/-. Grey shading in A and D denotes the dark photoperiod. Grey shading in C and F denotes 95% confidence intervals.

**Figure 6:**
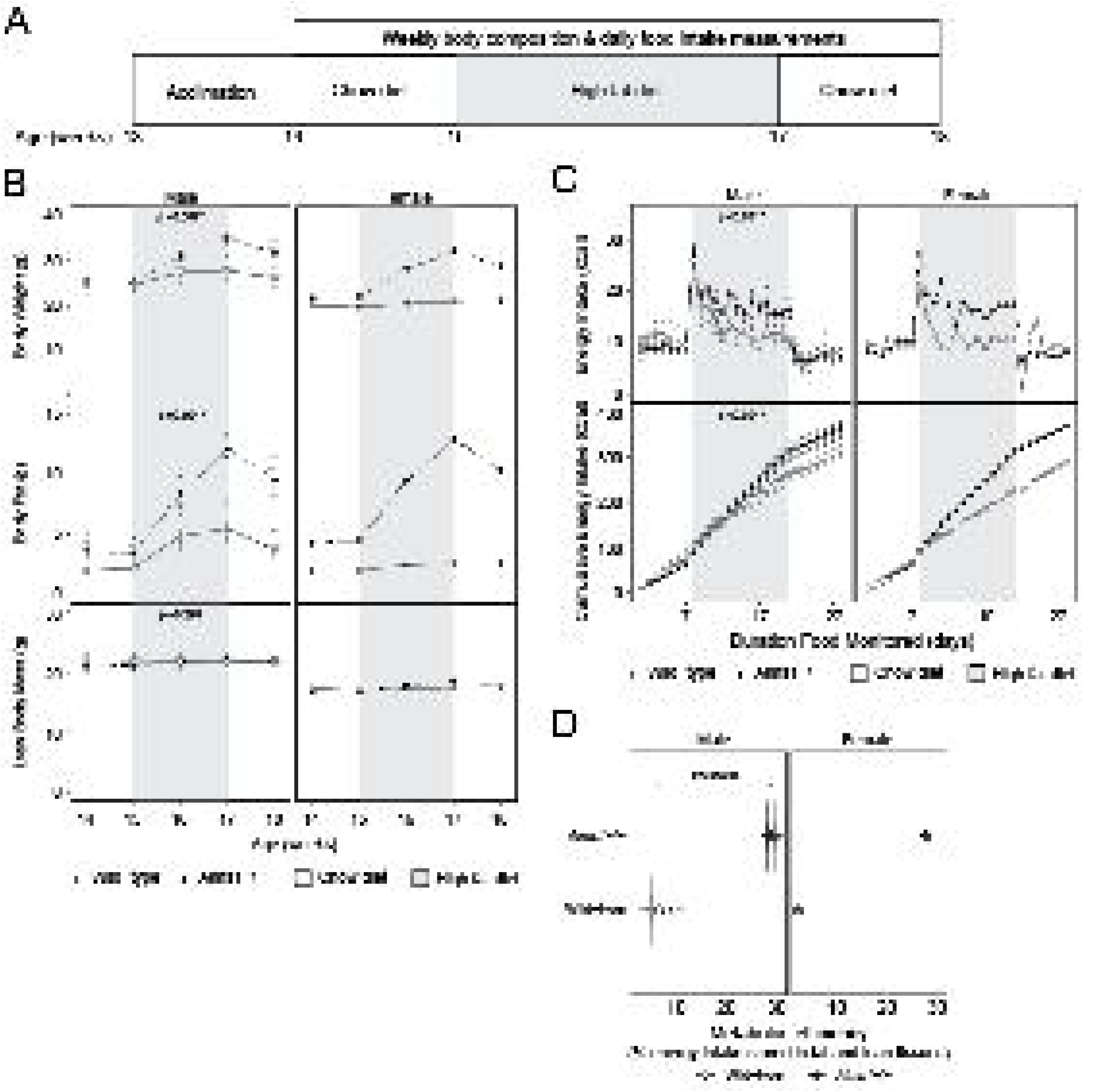
*Alms1*-/- mice eat more than WT mice when provided a palatable HFD. A. Overview of experimental design. B. Body composition of *Alms1*-/- and WT male (n=3 and n=4-5, respectively) and female mice (n=1 and n=1, respectively) before, during, and after a two-week period of high-fat feeding. C. Energy intake of *Alms1*-/- and WT male (n=3 and n=4-5, respectively) and female mice (n=1 and n=1, respectively) before, during, and after a two-week period of high-fat feeding. D. Metabolic efficiency during the high-fat feeding phase. Data presented are group means ± se with individual data overlayed (except for females where n=1/group). Open circles denote WT, closed circles denote *Alms1*-/-. Grey shading denotes HFD phase.

### Alms1-/- mice eat more than WT mice when provided a palatable HFD

Two previous studies reported that male and female *Alms1*-/- (*foz/foz*) mice consume more food than WT [16, 21]; however, we were unable to replicate this observation in our chow-fed *Alms1*-/- cohorts. To address this discrepancy, we sought to determine whether food palatability influenced food intake in a separate cohort of *Alms1*-/- mice (Figure 6). Mice were given one-week to acclimate to solo housing, after which chow intake was measured daily for one-week. Food was then switched from chow to a palatable HFD for two-weeks, and then back to chow for a final week. In addition to recording daily food intake over the four-week period, we measured body composition on a weekly basis (Figure 6A). In line with our earlier findings, compared to WT, *Alms1*-/- mice had increased adiposity while consuming chow (Figure 6B), yet did not consume more food than WT (Figure 6C). During the HFD phase, both WT and *Alms1*-/- gained bodyweight and increased in adiposity (Figure 6B); however, these gains were exacerbated in *Alms1*-/- (Figure 6B, upper panel, p=0.007, and center panel, p=0.001). Increases in bodyweight and adiposity in *Alms1*-/- appeared to be due to greater caloric intake compared to WT during this period (Figure 6C, p<0.001). Both WT and *Alms1*-/- lost weight and adiposity upon withdrawal of the HFD (Figure 6B), an observation associated with reduced energy intake due to the change in diet. No differences in energy intake were detected between genotypes after the diet was switched back to chow (Figure 6C). The metabolic efficiency of *Alms1-*/- mice increased compared to WT in response to the HFD (Figure 6D; p=3.5^-04^), suggesting that in the *Alms1*^*tvrm102*^ model of AS, increased energy intake only occurs in the setting of a palatable, high-energy diet and such a diet increases the ability of *Alms1*-/- mice to convert consumed energy to stored energy.

## Discussion

We performed comprehensive energy balance phenotyping of a mouse model of AS and demonstrate that in the *Alms1*^*tvrm102*^ model of AS, adiposity gains occur early and rapidly in male mice, but much later in females. Rapid increases in body fat in males are, in part, due to a marked reduction in EE during early life. Energy intake increase in a genotype-specific manner when mice are provided a palatable, high-energy diet, although chow data demonstrate that HFD-driven increases in food intake are not necessary for the initial establishment of obesity in this model. Interestingly, the EE deficit observed in male *Alms1*-/- mice does not persist as mice age, suggesting loss of Alms1 either causes a developmental delay in the mechanisms controlling EE, or that there is activation of compensatory mechanisms after obesity is established. Teasing out how Alms1/ALMS1 modulates EE and how sex moderates this process will be a key step toward understanding how mutations in *ALMS1* lead to childhood obesity in AS.

A cardinal feature of AS is rapid onset of obesity during early childhood [28]. The average body fat of individuals diagnosed with AS ranges from 34.7-38.4% [11, 34], with male patients thought to gain proportionally more body fat than females [34]. However, since patient cohort data are typically not separated by sex due to the ultra-rare incidence of AS [35], it has been challenging to parse out whether sex moderates the progression of obesity in patients. Animal models of AS should be able to fill this gap, although to date, body composition data collected for either sex has primarily been limited to single (or random) timepoints, making it difficult to infer anything about obesity onset and its progression over time [12, 15, 16]. We provide new insight into the timing of obesity onset and rate of obesity progression, demonstrating *Alms1*-/- males become obese earlier and gain adiposity more rapidly than females, supporting the preliminary observation that male patients with AS may gain more absolute body fat than female patients [34].

It has been suggested patients with AS develop obesity due to increased caloric intake as opposed to a deficit in resting EE [11]; however, data are inconclusive whether loss of ALMS1 function drives hyperphagia, hedonic intake, or if other determinants of energy balance also contribute to obesity in these patients [36]. Caregiver hyperphagia questionnaire scores suggest hyperphagia may be elevated in people with AS compared to age-, sex-, race-, and BMI-Z-score-matched control subjects [11], and short-term food intake measurements in the *foz/foz* model support this notion [16]. Besides these reports, limited data exist – in either animal models or patients – to support the assertion that obesity in AS is driven solely by overeating. To address this issue and strengthen our understanding of how mutations in *ALMS1*/*Alms1* disrupt energy balance, we performed comprehensive energy balance phenotyping in younger (8-week-old) and older (18-week-old) adult mice. EE was determined while simultaneously measuring food intake and physical activity in order to get a complete picture of how Alms1 influences the key physiologic mechanisms that determine energy balance. Unlike previous studies that suggest loss of Alms1 results in hyperphagia [16, 21], we did not observe any differences in food intake in *Alms1-/-* mice compared to WT mice. If anything, older male *Alms1*-/- mice tended to eat less when provided a chow diet. We did, however, find that when provided a palatable, high-energy diet, *Alms1*-/- mice ate more than WT mice, an effect completely reversed after returning the mice to a chow diet. This observation, together with the finding that pair-feeding *foz/foz* males (HFD matched to WT intake) does not prevent weight gain [21], suggests although some combination of food environment and an altered hedonic response to food might exacerbate obesity in *Alms1*-/- mice, it is not the primary driver.

Our observation that younger male mice have reduced EE compared to WT littermates further supports the notion that altered thermogenic capacity at an early age, rather than overeating, drives the obesity phenotype of *Alms1*-/- mice. In line with this, it has been reported that *foz/foz* males have comparable normalized VO_2_ (reported as VO_2_ per unit body mass scaled 2/3) when receiving a chow diet but fail to increase VO_2_ by the same magnitude as WT mice when consuming a HFD [21]. This suggests *foz/foz* mice have a defect in the mechanisms controlling the non-resting thermogenic pathways. Indeed, *foz/foz* mice are unable to maintain core body temperature when subjected to acute cold exposure, in part because both glucose uptake and induction of thermogenic transcripts is attenuated in brown adipose tissue [21]. Interestingly, repeated cold exposure (2 h/d at 4°C for 4-weeks) corrects the ability of HFD-fed *foz/foz* mice to maintain core temperature during a cold challenge, while also restoring the thermogenic profile of adipose tissues and slowing weight gain [21]. These observations suggest although Alms1 influences the bioenergetic response to acute thermogenic challenges, some deleterious effects caused by Alms1-loss can be overcome by ‘training’ the thermogenic pathways to respond. Here, reduced EE in male *Alms1*-/- mice could be explained by younger mice being less active and having reduced activity EE. Given the adaptability of cold tolerance previously observed in the *foz/foz* strain [21], it is plausible that increasing activity through regular exercise training may also help overcome the early life EE deficits of male *Alms1*-/- mice. To date, few studies have quantified activity levels in *Alms1*-/- mouse models, and no reports of physical activity in patients with AS are available. Given resting EE no longer appears to differ between patients with AS and age-, sex-, race-, and BMI Z-score-matched control subjects after adjusting for differences in lean mass [11], understanding how ALMS1 affects adaptive thermogenic pathways (exercise, NEAT, thermic effect of food) will be an important line of investigation that may open doors to new therapeutics aimed at increasing EE and attenuating weight gain.

Curiously, adiposity often reduces after patients with AS reach adulthood; however, the cause of this is debated [33, 35]. It has been suggested that the attenuation of obesity coincides with the onset or increased severity of other clinical complications [12], that weight loss in adults may be due to lifestyle modifications made after diabetes onset [33], or that adipose tissue becomes metabolically inflexible and a type of lipodystrophy develops [36]. In this study we did not follow the mice long enough to observe attenuation of adiposity gains with increasing age; however, we can speculate that our observation of age-associated reversal of the EE deficit seen in younger male mice via an increase in resting energy expenditure may contribute to this phenomenon. Increased EE without a change in energy intake would likely lead to slowing of weight gain and eventual weight loss, which might explain adiposity reductions observed in patients with AS after they reach adulthood. How or why EE deficits reverse over time in *Alms1*-/- mice and whether this phenomenon also occurs in patients with AS are important questions that future studies should consider.

## Conclusion

We strengthen our understanding of the role of ALMS1/Alms1 in energy balance regulation by demonstrating that *Alms1* mutations cause early and rapid adiposity gains in male mice, with adiposity gains occurring much later in females. Rapid increases in body fat in males were, in part, due to a reduction in early life EE and not increased energy intake in the setting of a chow diet. Increased energy intake did occur with exposure to a palatable, high-energy diet and did exacerbate adiposity gains in *Alms1*-/- mice; however, overeating was not required for obesity progression. If we were to extrapolate these findings to patients with AS, this suggests biological sex and food environment are major determinants of the rate at which obesity manifests in AS and its overall severity. Whether deficits in early life EE can be targeted to attenuate obesity severity in AS is an important question future studies should address.

## Acknowledgements

The authors wish to thank past and present members of the Han Lab, the Stephenson Lab, and our former colleagues at UTHSC for insightful discussions about this work.

