## Supplementary figures and images for "Energy expenditure deficits drive obesity in a mouse model of Alström syndrome"

### Supplementary Figure 1

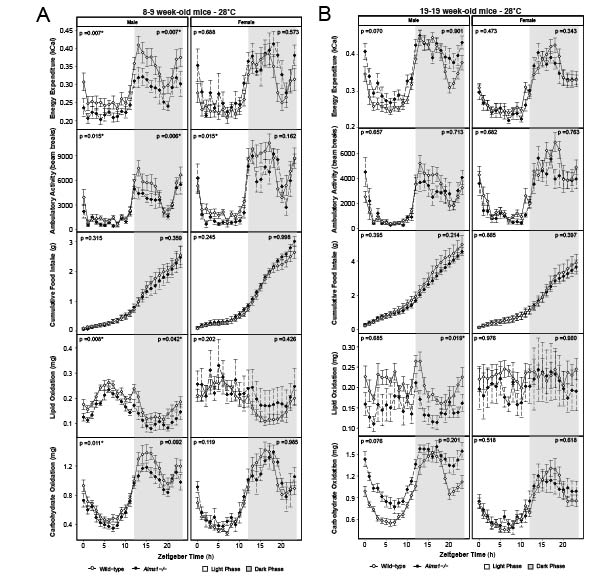
